# Circuits in the Motor Cortex Explain Oscillatory Responses to Transcranial Magnetic Stimulation

**DOI:** 10.1101/2023.07.04.547732

**Authors:** Lysea Haggie, Thor Besier, Angus McMorland

## Abstract

Transcranial Magnetic Stimulation (TMS) is a popular method used to investigate brain function. Stimulation over the motor cortex evokes muscle contractions known as motor evoked potentials (MEPs) and also high frequency volleys of electrical activity measured in the cervical spinal cord. The physiological mechanisms of these experimentally derived responses remain unclear, but it is thought that the connections between circuits of excitatory and inhibitory neurons play a vital role. Using a spiking neural network model of the motor cortex, we explained the generation of waves of activity, so called ‘I-waves’, following cortical stimulation. The model reproduces a number of experimentally known responses including direction of TMS, increased inhibition and changes in strength. Using populations of thousands of neurons in a model of cortical circuitry we showed that the cortex generated transient oscillatory responses without any tuning, and that neuron parameters such as refractory period and delays influenced the pattern and timing of those oscillations. By comparing our network with simpler, previously proposed circuits, we explored the contributions of specific connections and found that recurrent inhibitory connections are vital in producing later waves which significantly impact the production of motor evoked potentials in downstream muscles (Thickbroom, 2011). This model builds on previous work to increase our understanding of how complex circuitry of the cortex is involved in the generation of I-waves.

## Introduction

The activity of motor cortical neurons directly contributes to the generation and control of movement in humans. Non-invasive brain stimulation techniques, notably transcranial magnetic stimulation (TMS) have been used to study the connection from cortex to muscle in recent decades (Volz et al., 2015; Ni, Müller-Dahlhaus, et al., 2011). A single-pulse TMS at a sufficient intensity applied over the motor cortex evokes electrical currents in the neurons and a subsequent contraction in downstream muscles. Following the application of TMS over the motor cortex, high frequency volleys of electrical activity of 600–700 Hz have been recorded from the epidural space in the cervical spinal cord of humans, as well as cats and monkeys (Ziemann and Rothwell, 2000; Di Lazzaro, Ranieri, et al., 2012). These cortically-evoked responses in the pyramidal tracts were first described in Patton and Amassian (1954)’s study; however, the exact physiological mechanisms involved in the generation of these waves remain unclear.

Cortically-evoked volleys of electrical activity have been shown to be due to the direct and indirect activation of corticomotor cells. The initial volley, known as the D (direct)-wave, is considered to be due to the direct activation of the layer 5 pyramidal tract neurons. Following the D-wave, a series of three to four I (indirect)-waves have been attributed to the activation of upstream cortical neurons with differing synaptic delays in recurrent or polysynaptic connections with the pyramidal neurons (Di Lazzaro, Ranieri, et al., 2012; Rusu et al., 2014). I-waves originate from motor cortex circuits and are susceptible to cortical injuries, unlike the D-wave which is relatively independent of the functional state of the cortex (Zewdie and Kirton, 2016; Patton and Amassian, 1954). Hypothetical models have been proposed to explain I-wave generation involving neuronal circuits of excitatory and inhibitory interneurons, but these have not been explicitly tested in populationbased models (Ziemann and Rothwell, 2000; Di Lazzaro and Ziemann, 2013).

The motor cortex is composed of a large number of neurons with different size, location, orientation and functions (Young, Collins, and Kaas, 2013). The neurons of the motor cortex are structured in a highly stereotyped manner of layers and columns (Douglas and Martin, 2004; Hatsopoulos, Xu, and Amit, 2007). Canonical cortical circuits of excitatory and inhibitory neurons have previously been suggested by Gilbert and Wiesel (1983) and Douglas and Martin (2004), involving recurrent connections between the six layers of the cortex. The circuits in the motor cortex and how they are activated by TMS is difficult to establish through anatomical and electrophysiological measurement alone because it is currently infeasible to record from and manipulate precisely the large numbers of neurons involved. Though there have been some recent efforts to experimentally explore the cellular mechanisms of TMS, computational modelling can test the plausibility of proposed mechanisms and shed light on hypotheses involving complex interactions between larger-scale neuron groups (Epstein, 2008; Murphy et al., 2016).

The generation of I-waves has previously been modelled using compartmental models of individual neurons and spiking neural networks. Rusu et al. (2014) modelled a population of layer 2 and 3 neurons projecting onto a detailed layer 5 pyramidal cell model and suggested that I-waves are generated by the delay due to synaptic inputs arriving at different parts of the layer 5 dendritic tree. This was a feedforward model and ignored any recurrent connectivity which exists within and between cortical layers (Weiler et al., 2008). Esser, Hill, and Tononi (2005) used a large-scale spiking neural network with detailed anatomical and physiological constraints to model the motor cortex and the effects of *GABA*_*A*_ enhancement, as well as single and paired-pulse TMS. However, this study has not been replicated and did not identify or examine the key contributions of the components of the circuit, rather it treated the whole neural network as a black box without showing which connections or features were important in capturing the TMS response.

Activation of TMS-induced descending volleys is dependent on the orientation of the TMS coil due to the direction of induced current (Ni, Charab, et al., 2011). A current in the posterior-anterior (PA) direction preferentially activates early I-waves which increase in amplitude and number as stimulation intensity increases. Stimulation in the anterior-posterior (AP) current direction results in fewer, lower amplitude waves which have a longer onset latency. Multiscale computational modelling exploring the effects of TMS on individual neurons suggests that TMS activates neuronal axons aligned to the direction of the induced electric field, depending on the cortical morphology (Aberra et al., 2020; Seo et al., 2017). Shifting the direction of TMS from an AP to PA direction potentially activates different circuits of the motor cortex. The later onset of AP-stimulation-induced I-waves is thought to be the result of polysynaptic, pathways through the premotor cortex (Spampinato, 2020).

While previous computational models have captured several properties of I-waves, a coherent conceptual framework for understanding the response of cortical circuits to TMS is lacking. The aim of this study was to develop a functional model of the primary motor cortical circuitry and determine whether recurrent connections in the cortex explain physiological responses to TMS as observed by the generation of I-waves in the spinal column. We used a spiking neural network model of excitatory and inhibitory neurons in the different cortical layers, connected with plausible connection probabilities and synaptic strengths. This motor cortex model is described in Haggie, Besier, and McMorland (2022) and uses elements from a previously published cortical model by Potjans and Diesmann (2014) and the model put forth by Esser, Hill, and Tononi (2005).

The model was altered in several ways to explore the contributions of the structural arrangement and functional connectivity between neuron groups to the response of TMS over a region of motor cortex, including testing several circuits previously hypothesised by Di Lazzaro, Ranieri, et al. (2012) to be required for I-wave generation. Our model replicates the effects of single pulse PA and AP TMS, increased GABA-ergic inhibition, and facilitatory paired-pulse TMS and also shows adaptation to TMS strength and a cortical silent period. To the best of our knowledge, this is the first study to use a large-scale population model of the response to TMS in the AP and PA current directions explaining circuit-level contributions.

## Methods

The motor cortex model used in this simulation is a spiking neural network containing 38,556 leaky integrate-and-fire neurons and over 160 million synaptic connections characterising the laminar structure of the cortex. Eight neuron groups represents excitatory and inhibitory neurons in layers 2/3, 4, 5 and 6. The leaky-integrate and fire neuron model (equations 1, 2 and 3), and connectivity were adapted from previous work by Potjans and Diesmann (2014) and Esser, Hill, and Tononi (2005). Connectivity is locally defined by a Gaussian probability based on spatial distances described by equation 4. Parameters for this model are shown in table 1 and the connections are described in more detail in Haggie, Besier, and McMorland (2022). This model portrays 1 mm^2^ of surface area of the motor cortex as shown in figure 1 and the connectivity between groups is defined by various delays and weightings represented by the circuit diagram in figure 2.

**Table 1.** Parameters for neuron model. Taken from Potjans and Diesmann (2014).

| Symbol | Name | Value |
| --- | --- | --- |
| $C_m$ | Membrane Capacitance | 250 pF |
| $\theta$ | Threshold | -50 mV |
| $\tau_{ref}$ | Refractory Time | 2 ms |
| $\tau_m$ | Membrane Time Constant | 10 ms |
| $\tau_{syn}$ | Synaptic Time Constant | 0.5 ms |
| $V_r$ | Reset Value | -65 mV |
| $g_e$ | Excitatory Conductance | 1 |
| $g_i$ | Inhibitory Conductance | -4 |
| $w$ | Synaptic Weight | 87.8pA |
| $d_e$ | Mean Excitatory Delay | 1.5 ms |
| $d_i$ | Mean Inhibitory Delay | 0.8 ms |

**Figure 1.**
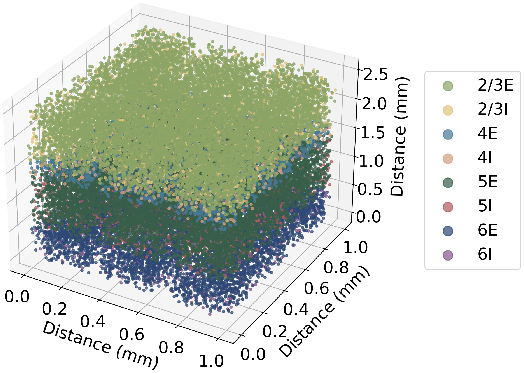
Spatial arrangement of neurons in the cortical model showing the laminar structure, representing a 1*mm*^2^ surface of cortex with a thickness of 2.3 mm.

**Figure 2.**
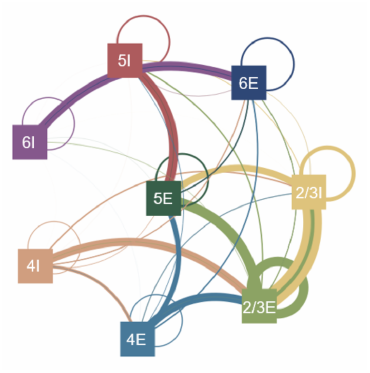
Circuit diagram of model showing connections between the populations of excitatory (E) and inhibitory (I) neuron groups. Groups are organised according to their layer. The colour of the lines indicates the source group. Thickness is indicative of relative number of connections and weighting of synapses.

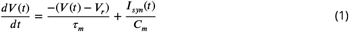

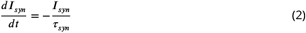

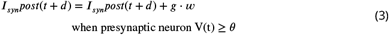

when presynaptic neuron V(t) *≥ θ*

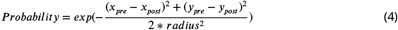

A consistent connectivity was defined in the TMS simulation studies by reading in tables of preand post-synaptic neuron indices corresponding to a single instance of the cortical model with local connectivity. A single instance of the model was used in most simulations to eliminate the possibility that differences observed were due to specific details of the randomly chosen network structure. We observed similar results in the network firing rates and patterns as well as observed responses to a simulated TMS perturbation, even with slight changes to the network structure based on the randomness of input, spatial and connectivity definition in this model.

External input to the model representing activity received by motor cortex neurons from other brain regions was a Poisson-distributed spike train of 8 Hz. The number of these inputs for each neuron was constant at 2000 inputs for excitatory inputs and 1850 for inhibitory inputs, as previously used in the layer-independent protocol from Potjans and Diesmann (2014). The spontaneous activity of the model shows a resting state population firing rate of 3.2 Hz in layer 2/3E and 10.5 Hz in layer 5E neuron groups of the cortex which is similar to intracortical recordings (Dabrowska et al., 2021). Figure 3 shows the spontaneous activity of the layer 2/3E and layer 5E neuron populations in which bands of oscillatory waves of activity can be observed similar to those reported in Esser, Hill, and Tononi (2005).

**Figure 3.**
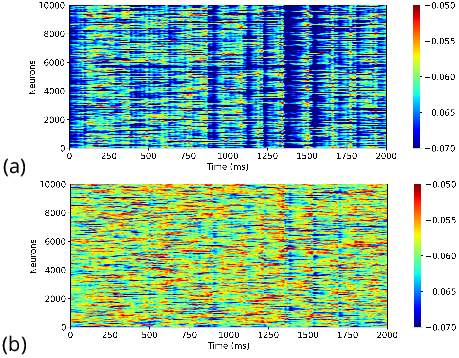
Spontaneous firing activity in (a) layer 2/3E and (b) layer 5E neurons of the motor cortex model with a random, layer-independent input of 8 Hz firing from a Poisson distribution.

TMS has been shown to induce the activation of single neurons in areas of the cortex measuring less than 2 mm in diameter (Romero et al., 2019). The effect of TMS is also thought to act superficially on the cortex with reduced strength at further depths (Rudiak and Marg, 1994; Trillenberg et al., 2012). A single TMS pulse was modelled as a direct input current to a proportion of individual neurons, i.e. a value of 25% of fibre terminals stimulated, which is the proportion of fibres stimulated in non-human primate studies to elicit motor evoked potentials (Romero et al., 2019). The strength of the TMS activation at different depths was modelled with a linear reduction in the input current to neurons as the depth (Z) increases (figure 4). Different amplitudes of TMS were modelled by altering the proportion of neurons activated.

**Figure 4.**
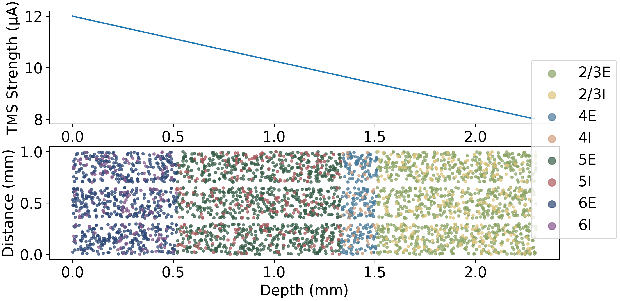
Model simulation of TMS as an input current to neurons where the strength in the Z direction linearly decreases with depth.

A group of neurons representing the premotor groups was added to the cortical model to explore the different pathways activated during AP and PA TMS protocols. The majority of neurons from the premotor area projects to layer 2/3 in the motor cortex with a small proportion of connections to layer 5 (Lu, Preston, and Strick, 1994; Matsumura and Kubota, 1979; Leichnetz, 1986; Ghosh, Brinkman, and Porter, 1987). The response to PA TMS was modelled by the activation of the primary motor cortical neurons while the AP response to TMS was modelled by stimulation of the premotor neuron group, shown in figure 5. Previous computational models have suggested that TMS activates axonal collaterals aligned to local electric field direction and that AP stimulation results in activation of collaterals directed toward the premotor cortex, resulting in fewer and delayed I-waves (De Geeter et al., 2015; Aberra et al., 2020).

**Figure 5.**
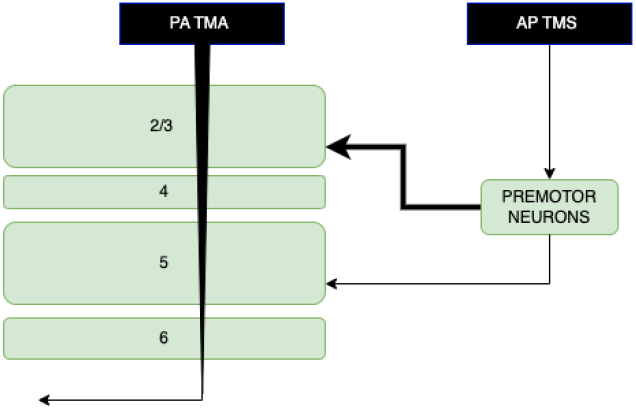
Diagram showing circuits of the cortex involved in the generation of I-waves in response to Transcranial Magnetic Stimulation (TMS) in the Anterior-Posterior (AP) and Posterior-Anterior (PA) directions.

Di Lazzaro, Ranieri, et al. (2012) has previously described a minimal cortical circuit hypothesised to generate I-waves. Components of this circuit were tested using the motor cortex model to establish the contributions to synchronised wave generation. A diagram of the proposed circuits by Di Lazzaro, Ranieri, et al. (2012) are shown in figure 6 below. To create these simpler circuits, the same neuron indices were used for the populations of layer 2/3E, 2/3I and 5E and only the additional neuron groups and connections were removed so the network in the relevant populations were identical to the larger network model.

**Figure 6.**
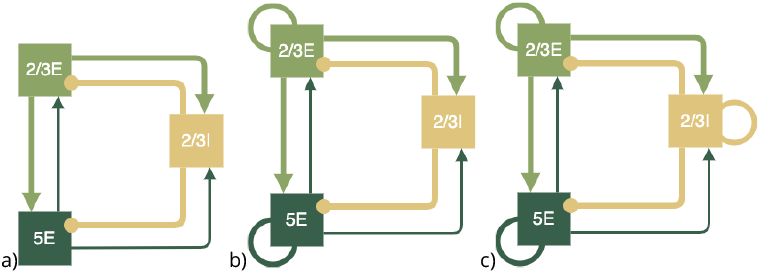
Simpler cortical circuits with key neuron groups of layer 2/3 and 5 hypothesised to be the key neurons involved in the generation of I-waves, as suggested by Di Lazzaro, Ranieri, et al. (2012)

All differential equations were solved using a 4th order Runge-Kutta numerical integration with a step size of 0.1 ms. A total of 250 ms of neuronal activity was simulated and the last 200 ms of steady state model behaviour was analysed. The TMS activation was applied 150 ms into the simulation. The model took 5 minutes to run on a six core CPU. Firing frequencies were calculated from the instantaneous spiking activity of the entire population in each time window and smoothed using a Gaussian weighted average with a standard deviation of 0.15 ms.

The spiking frequency of neurons during each simulation was recorded and I-waves were interpreted to be the cumulative activity of layer 5 excitatory neurons. The axons of layer 5 excitatory neurons made up the majority of the corticospinal tract where I-waves are physiologically recorded.

## Results

The model replicated the high frequency activity observed in layer 5 pyramidal tract neurons in response to a simulated TMS. I-wave activity has been recorded from the epidural space of the cervical spinal cord in humans following simulation of the motor cortex. The measured changes in electrical potential were assumed to be the summation of the action potentials in the axons of layer 5 neurons which extend down from the motor cortex (Esser, Hill, and Tononi, 2005; Rusu et al., 2014). From the simulation, a D-wave at the time of TMS delivery and series of three I-waves of neuron activity with interspike intervals (ISIs) of 1.2–1.5 ms was observed. These results were similar to the response recorded in humans, is shown in figure 7a. A delayed two-spike response with lower amplitudes in the model was observed in response to simulated AP TMS as shown in figure 7b.

**Figure 7.**
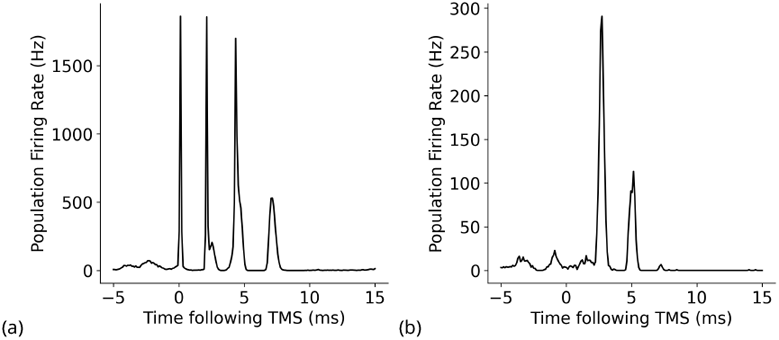
Cortical response in the model’s layer 5E neurons to (a) PA TMS stimulation and (b) AP TMS stimulation

Figure 8 shows different numbers of waves appear in each layer of the cortical model during PA stimulation. This model suggests that all cortical layers are involved in the generation of Iwaves, but further analysis discussed later in this section suggests that particular groups may be key contributors. A raster plot of neuron firing arranged by firing frequency is shown in figure 9. This plot shows that neurons contribute to the firing behaviour of the population unequally, a result supported by experimental measures of neuron firing in the cortex (Dabrowska et al., 2021; Tomov et al., 2014; Lacey et al., 2014; Roxin et al., 2011).

**Figure 8.**
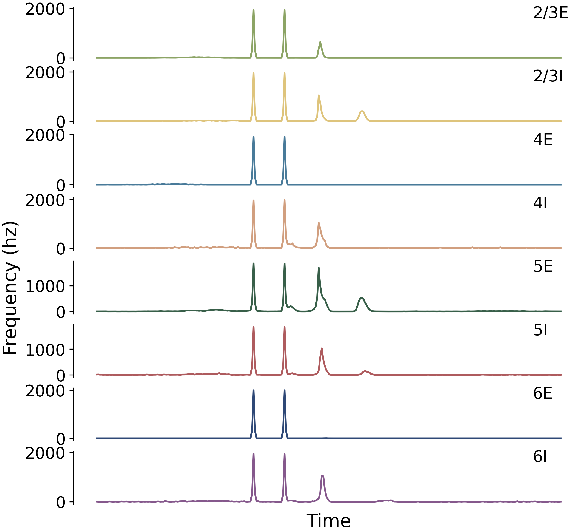
The response in firing activity to TMS in each neuron group of the model.

**Figure 9.**
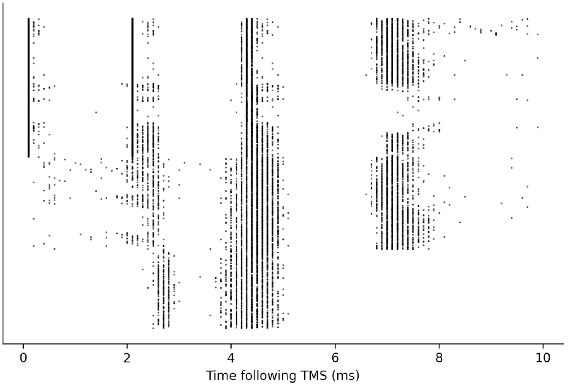
Raster plot of layer 5E neurons arranged by number of firing instances.

A cortical silent period was observed immediately after an applied TMS, similar to those seen experimentally (figure 10). The silent period is described as a pause in electromyographic activity following the onset of a motor evoked potential in response to TMS during voluntary muscle contraction and is thought to be mediated by both cortical and spinal mechanisms though the contributions of each are still unclear. Complete loss of the silent period has been observed in patients with ischemic lesions in the motor cortex suggesting that the silent period is of cortical origin (Schnitzler and Benecke, 1994). Silent period duration shows large individual variation but typically ranges from 100–300 ms with duration increasing as TMS strength increases (Ahonen et al., 1998). Figure 10 shows a period of approximately 50 ms with a substantially decreased level of spiking activity following TMS, particularly in excitatory neurons, which supports the theory that cortical mechanisms may contribute to the silent period (Schnitzler and Benecke, 1994; Özyurt et al., 2019).

**Figure 10.**
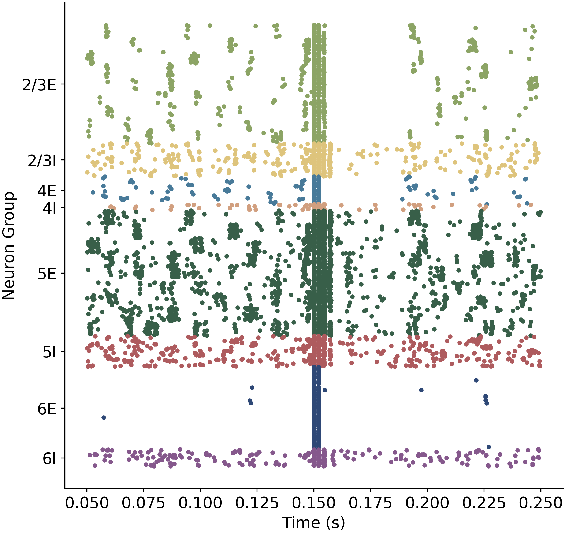
Raster plot of all cortical neurons with cortical silent period in layers 2/3E and 4E observed after the TMS simulation at 0.150 s.

In the PA direction, the strength of TMS affected the amplitude of the resulting I-waves. Figure 11 shows the amplitude of waves in response to increasing the strength of PA TMS, showing a similar trend to that recorded by Nakamura et al. (1996). Increasing the strength of the TMS, by increasing the proportion of neurons stimulated, increases the resulting amplitude of the population frequency response. The I3 wave was recruited at a higher TMS strength in this model, with the I3 wave only appearing when 22.5% of the neuron population was stimulated. This was similar to experimental results for the proportion of neurons affected by single pulse TMS to evoke an effect showing burst of action potentials in neurons recorded by Romero et al. (2019). The I3 wave is also thought to play a disproportionately significant role in the recruitment of motoneurons and size of the MEP (Thickbroom, 2011).

**Figure 11.**
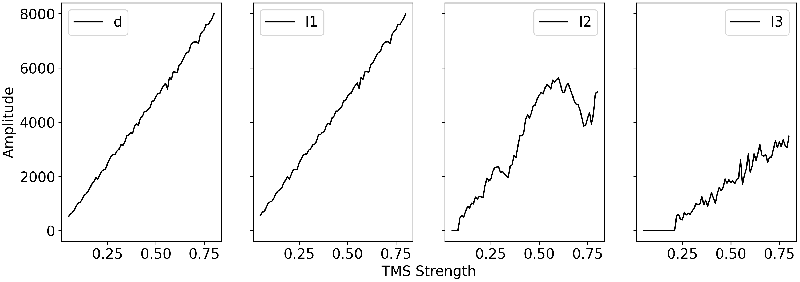
Plots of I-wave amplitudes in response to changes in TMS strength. Amplitudes increase with increased strength of TMS, similar to experimental results presented by Nakamura et al. (1996).

An increase in the strength of inhibitory connections (by 1.4 times) to simulate the effect of pharmacological GABA_A_ agonists resulted in an observed decrease in the amplitude of I2 and I3. Figure 12 shows the population based firing activity following TMS when the strength of inhibitory connections was increased. This finding corresponds to observations in pharmacological experiments with TMS, where administration of a benzodiazepine such as lorazepam suppresses later I-waves but not the first I-wave (Di Lazzaro, Ranieri, et al., 2012). Similarly, I1 waves are rarely affected by conditioning stimuli or enhancement of GABAergic activity (Zewdie and Kirton, 2016; Di Lazzaro and Ziemann, 2013).

**Figure 12.**
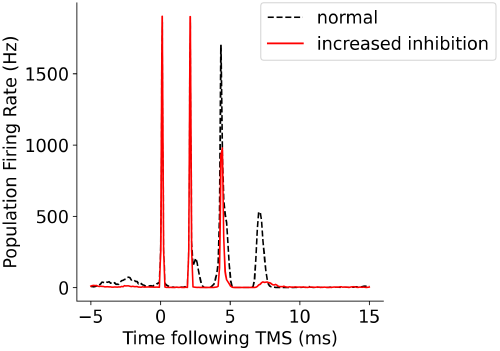
Effect of increased inhibition on later I-waves showing suppression of later I-waves (I2 and I3) as observed in experimental results (Hicks et al., 1992; Burke et al., 1993; Woodforth et al., 1999).

**Figure 13.**
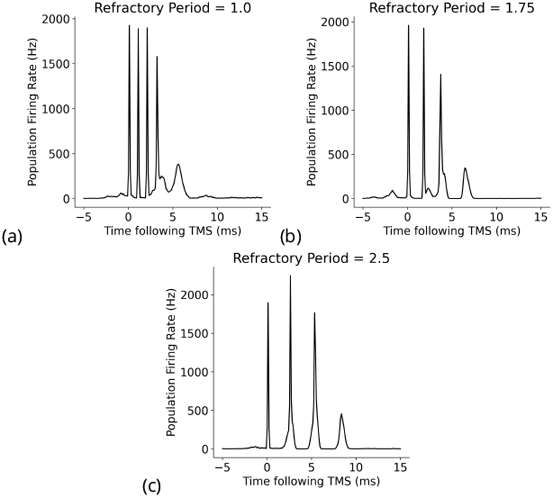
Effect of changing the refractory period of neurons on I-waves from the original value of 2.0 ms, where (a) refractory period = 1.0 ms, (b) refractory period = 1.75 ms, (c) refractory period = 2.5 ms.

**Figure 14.**
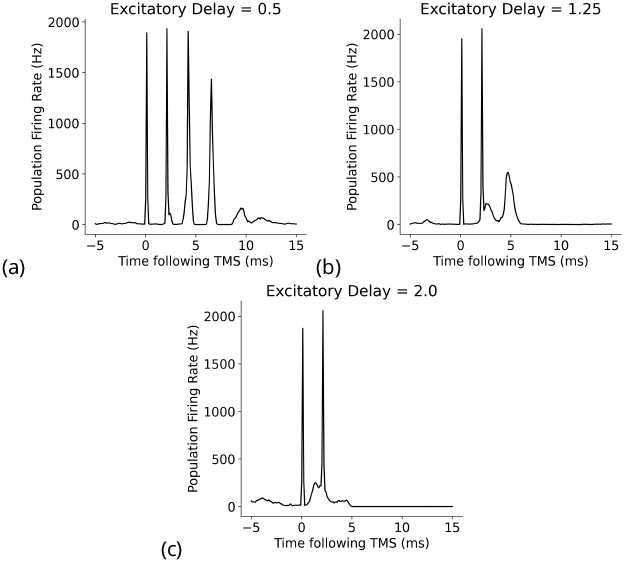
Effect of changes to the mean excitatory delay on I-waves from the original value of 1.5 ms, where (a) mean excitatory delay = 0.5 ms, (b) mean excitatory delay = 1.25 ms and (c) mean excitatory delay = 2.0 ms.

**Figure 15.**
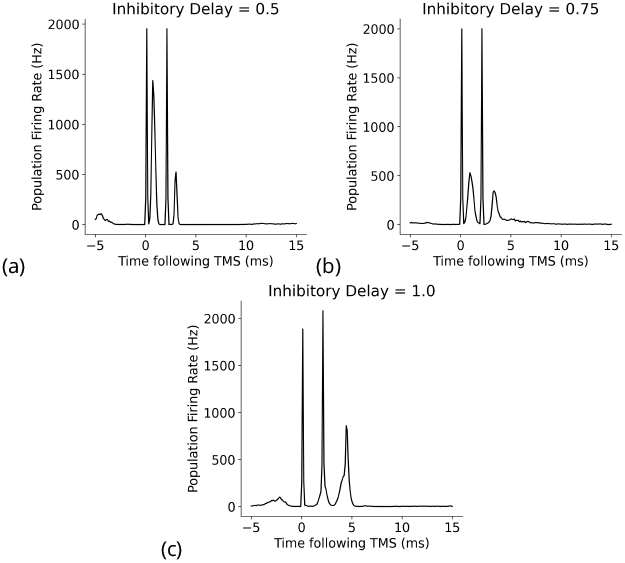
Effect of changes to the mean inhibitory delay on I-waves from the original value of 0.8 ms, where (a) mean inhibitory delay = 0.5 ms, (b) mean inhibitory delay = 0.75 ms and (c) mean inhibitory delay = 1.0 ms.

Next, we assessed the relationships between the temporal parameters of individual neurons, refractory period, and transmission delay, and the amplitude and timing of I-waves. A decrease in refractory period led to a higher frequency response with the time between the waves decreasing from 2 ms at a refractory period of 1.5 ms, to 1.2 ms at a refractory period of 1 ms. Changes to excitatory and inhibitory delays had the biggest effect on later I-waves (ie. I2 and I3), but in opposing directions. Increasing the excitatory delay resulted in a left shift (earlier in time) of the later I-waves, whereas increasing the inhibitory delay resulted in a right shift (later in time) of the later I-waves. The amplitude of the D-wave remained constant, as expected, and I1 was also consistent except in some cases when the later I-waves may have superimposed, meaning activity of the I1 and I2 waves was combined, resulting in an increased amplitude.

Simplified circuits with specific components of the full model omitted were tested to investigate the contributions of different network components to the generation of I-waves. A minimal circuit which contained superficial layer 2/3 neurons and deep layer 5 neurons, as well as a group of inhibitory neurons based on previous models put forth by Di Lazzaro, Ranieri, et al. (2012) were tested. The initial circuit with no recurrent connections between the layers (figure 16) resulted only in an I1 wave being produced, suggesting that the direct connections between layer 2/3 and 5 are responsible for the generation of I1, and that more complex circuits with recurrent connections are necessary to produce later I waves.

**Figure 16.**
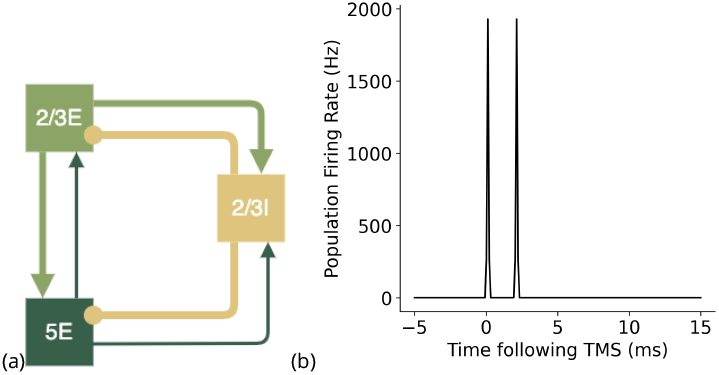
A simple cortical circuit with layer 2/3E, 2/3I and 5E neuron groups with only interneuron group connections and no recurrent connection, shown in (a), produces I1 wave, shown in (b).

I-waves have previously been hypothesised to be generated by recurrent cortical layer 2/3 neurons (Seo et al., 2017). However, the addition of recurrent connections to excitatory neuron groups without recurrent inhibition did not result in the generation of an I2 wave and only in combination with recurrent inhibitory connections were later I-waves produced (figure 17). Inhibitory recurrent connections are necessary for an I2 wave to be produced, in addition to the initial I1 wave (figure 18).

**Figure 17.**
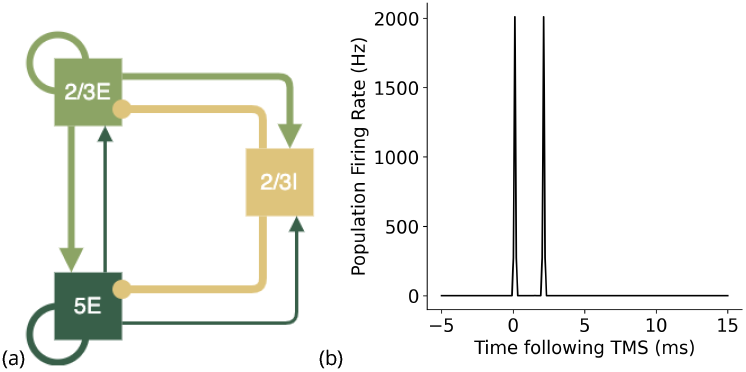
A simple cortical circuit (Di Lazzaro, Ranieri, et al., 2012) with excitatory recurrent connections in layer 2/3E and layer 5E neuron groups but no recurrent inhibitory connections, shown in (a), also only produces an I1 wave, shown in (b).

**Figure 18.**
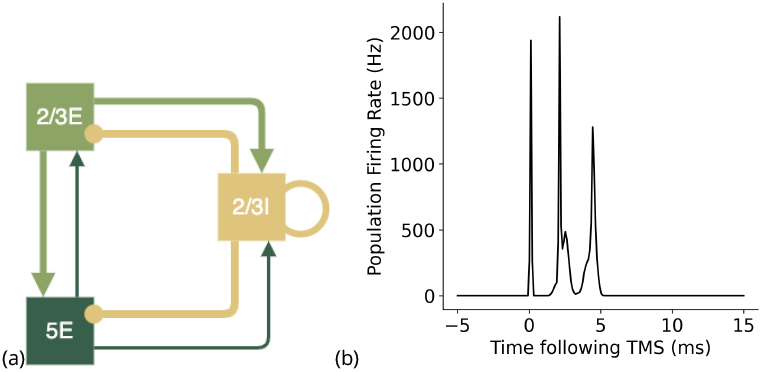
A simple cortical circuit (Di Lazzaro, Ranieri, et al., 2012) with recurrent inhibitory connections but no recurrent excitatory connections, shown in (a), produces both an I1 and I2 wave, shown in (b).

Recurrent inhibitory connections are vital for stable, balanced activity and the transient nature of the oscillations in response to stimulation. Increased excitation, through recurrent excitatory and inhibitory connections (which increase excitation and decrease inhibition in the neuron populations) such as in the circuit shown in figure 19a, resulted in a higher rate of firing in the network and continuous synchronised oscillations (figure 19b), in the model even in the absence of spontaneous input or stimulation.

**Figure 19.**
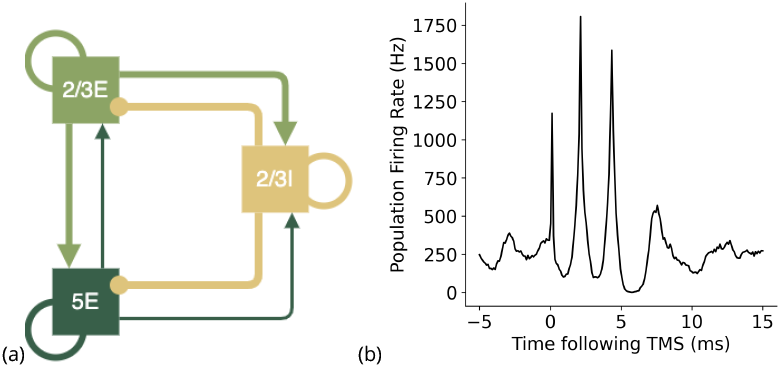
A simple cortical circuit (Di Lazzaro, Ranieri, et al., 2012) with recurrent connections in all neuron groups, shown in (a), produces both an I1 and I2 wave, but also generates continues high frequency oscillatory activity due to increased excitatory activity.

**Figure 20.**
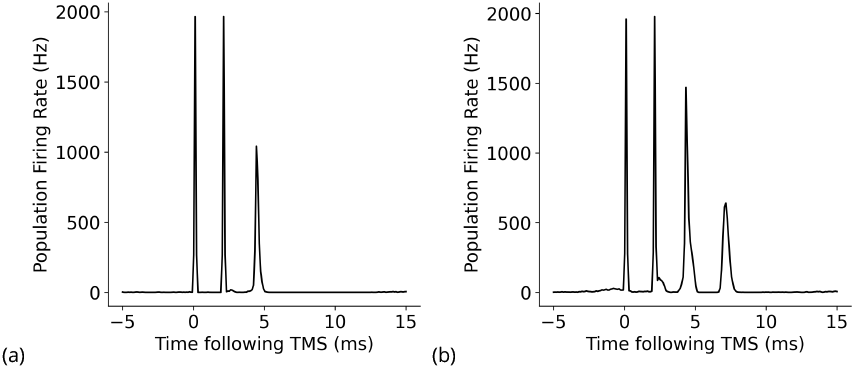
TMS response to circuit with a) no layer 4 excitatory or inhibitory connections, b) no layer 6 excitatory or inhibitory connections

Polysynaptic connections to and from other cortical layers such as 4 or 6 also contribute to the generation of later I-waves. Without layer 4 connections in the circuit, the model only generated I1 and I2; without layer 6 connections, the model generated three I-waves but with a lower amplitude I3 which, as mentioned previously, may be important for the recruitment of motor units and the generation of the MEP. These data suggest that while layers 2/3 and 5 are critical for I-wave generation, other cortical layers might also play a role in shaping the I-wave pattern.

The effect of paired pulse TMS on the motor evoked potential, usually measured at a distal muscle by electromyography (EMG), can be either facilitatory or inhibitory of motor evoked potentials depending on stimulus strength and the interval period (Müller-Dahlhaus, Liu, and Ziemann, 2008). Short Interval Intracortical Faciliation occurs when the first (conditioning) stimulus is, or both the conditioning and subsequent test stimulus are, administered at or above the motor threshold required to elicit motor evoked potentials, and stimuli are administered around 1.5 ms, 3 ms or 4.5 ms apart. Alignment of the interstimulus intervals with the timings of the I-wave peaks results in facilitatory interaction since the waves and stimuli are in-phase (Ziemann, Tergau, Wassermann, et al., 1998; Di Lazzaro, Ranieri, et al., 2012; Di Lazzaro and Ziemann, 2013).

Using our model of the motor cortex, paired pulse stimulation with an interstimulus interval of 2 ms, to match the interval of the I-waves, resulted in the increased amplitude of I1, I2 and I3 with an extra wave (ie. a small I4) (figure 21). Paired TMS at short intervals of 1.0 to 1.4 ms has been shown experimentally to be accompanied by larger and more numerous I-waves, with a more significant increase in I2 and I3 which matches our model results (Di Lazzaro, Rothwell, et al., 1999; Ziemann and Rothwell, 2000). The observed increased I-wave amplitudes in the model could contribute to explaining short interval facilitation observed in paired pulse protocols which increase the size of the resulting MEP when a second pulse is administered at specific time intervals due to the interaction of I-waves at the level of the motor cortex (Ziemann, Tergau, Wassermann, et al., 1998).

**Figure 21.**
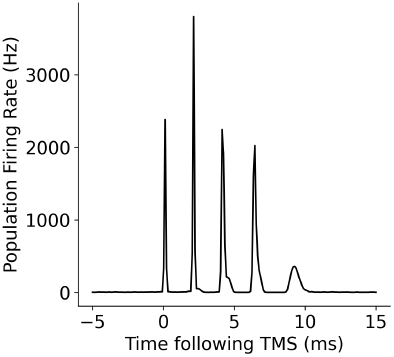
TMS response to paired stimulation at the same strength with an interval of 2ms.

## Discussion

A spiking neural network model of motor cortex circuitry has been developed and used to simulate and explain the cortical response of TMS on corticospinal neurons. The model replicates the inputoutput characteristics of single and paired pulse stimulation, as well as differences between PA and AP TMS, in regards to I-wave activity as synchronous high frequency network activity. The effects of TMS strength and increased inhibition were also captured by this cortical circuit model. The model incorporated excitatory and inhibitory populations of individual neurons, replicating the known physiological structure of the cortex including layers and columns. This circuit was also compared to simpler circuits previously proposed to be involved in the generation of I-waves (Di Lazzaro, Ranieri, et al., 2012). Connectivity between neurons was defined at the population level, and consequently the resulting model gives insight into how groups of neurons can interact to produce cortical responses observed experimentally.

Di Lazzaro, Ranieri, et al. (2012) hypothesised that the physiological generation of I-waves could be minimally represented by interactions between the circuits of inhibitory and excitatory neuron groups in layers 2/3 and 5 of the motor cortex. Our model uses simple LIF neurons situated in a large-scale cortical circuit based on previously published electrophysiological connectivity data to explain the origin of I waves. In addition to the full model, we tested simpler circuits representative of those proposed by Di Lazzaro, Ranieri, et al. (2012). The model demonstrated that I-waves can be explained by the recurrently connected circuitry of groups of neurons in the cortex. The results of this model support the idea that, while a subset of the connectivity is responsible for the generation of the initial I-waves, other cortical layers are likely involved in the generation of the full pattern of produced I-waves, rather than simply layer 2/3 and 5 which have been the main focus of previous work (Rusu et al., 2014; Di Lazzaro, Ranieri, et al., 2012).

The construction of this cortical model contained excitatory and inhibitory neuron groups in layers 2/3, 4, 5 and 6 with specific connections within and between each layer based on neuroanatomical data (Binzegger, Douglas, and Martin, 2004; Thomson et al., 2002). An important advancement of our model beyond that of Esser, Hill, and Tononi (2005) is the inclusion of a layer 4 neuron group, which has recently been shown to exist in the motor cortex (Yamawaki et al., 2014; Barbas and Garcia-Cabezas, 2015). Interestingly, the model reproduced three I-waves without the layer 6 excitatory and inhibitory neuron groups, but not without layer 4 neuron groups, which differs to the Esser, Hill, and Tononi (2005) model which represented layers 2/3, 5 and 6. The difference between the layers’ effect on the generation of I-waves may be due to the differing density of connections between the groups with more dense connections in layer 4. We took a similar modelling approach to Esser, Hill, and Tononi (2005) but used different neuron models and connectivity, though both connectivities were physiologically based. Both spiking neural network approaches were able to reproduce I-waves suggesting that recurrent, polysynaptic circuitry in the cortex is critical for generating synchronised oscillatory activity.

Paired pulse protocols can result in either the facilitation or the inhibition of the motor evoked potential Lazzaro et al., 1998. Epidural spinal cord recordings have shown larger and more numerous I-waves with paired-pulse TMS at short interstimulus intervals of 1.0–1.4 ms, demonstrating the intracortical origin of short-interval cortical facilitation (SICF), as replicated in this model (Di Lazzaro, Rothwell, et al., 1999). A subthreshold stimulus, delivered more than 6-25ms prior to a suprathreshold stimulus, resulted in facilitation whereas a subthreshold stimulus delivered less than 5 ms followed by a suprathreshold stimulus suppresses the MEP amplitude (**_sectionZewdie2016**; Peurala et al., 2008). Suppression of late I-waves has also been observed by direct epidural recordings suggesting a subthreshold stimulus may activate intracortical inhibitory circuits (Lazzaro et al., 1998). A subthreshold paired pulse stimulus activating inhibitory neurons could be explored further, with changes to the TMS and inhibitory neuron model. In addition, an inhibitory-specific neuron model and the resulting changes to network dynamics as well as I-wave generation could be future areas of investigation.

GABAergic interneurons play a key role in modulating excitatory activity within circuits of the cortex. GABA agonists have an effect on the amplitude and increased inhibition can suppress later I-waves, as demonstrated in this model (Ziemann, Tergau, Wischer, et al., 1998; Tremblay, Lee, and Rudy, 2016). This model also illustrated the importance of recurrent inhibitory connections in I-wave generation, particularly in the generation of I2. Inhibitory interneurons are thought to have important roles in cortical functions including modulation of cortical circuits through gain, regulation of pyramidal cell firing, plasticity, synchronisation and the generation of cortical patterns necessary for information transfer (Tremblay, Lee, and Rudy, 2016). This model shows that inhibitory connections are important in the generation of I-waves and corticomuscular transmission.

The differing results generated by testing variations in circuitry in this cortical model support the idea of separate functionality for distinct neuronal populations. Tsubo, Isomura, and Fukai (2013) suggested that the layers work as functional units with layer 2/3, serving as a “resonant oscillator” and layer 5 pyramidal neurons operating as “integrators”. Other hypothesised mechanisms of Iwave generation include the individual pyramidal cells of layer 5 operating as neural oscillators, repetitively discharging in response to depolarisation with high frequency bursting due to their intrinsic membrane and geometric properties (Triesch, Zrenner, and Ziemann, 2015; Silva, Amitai, and Connors, 1991; Kernell and Chien-Pingt, 1967; Hedrick and Waters, 2011; Kock et al., 2021). Neurons anatomically span several layers and are typically organised into layers based on the location of their cell body (Larkum et al., 2018). Although the laminar organisation in the cortex was preserved in this model, the heterogeneity of neurons in regards to morphology and electrophysiological behaviour even within each layer cannot be understated (Oswald et al., 2013). Specifically in the motor cortex, layer 5 has previously been separated into layer 5a and 5b due to the cells being larger and more numerous in the lower section of layer 5 (Sherwood and Hof, 2007).

Our representation of homogeneous populations of single compartment neurons in excitatory and inhibitory neuron groups cannot assess the consequence of neuron morphology, which plays a role in the integration of input at the neuron level (Seo et al., 2017). More detailed compartment models of layer 5 pyramidal neurons have been previously developed Rusu et al. (2014), whereas our work aims to describe possible roles of connectivity and neural population dynamics in the generation of I-waves and, for practical computational reasons, benefited from modelling simple point neurons. The complexity of simulating many thousands of multi-unit neuron models is still limited by computational power. Previous models have also explored complex neurotransmitter and synaptic models (Esser, Hill, and Tononi, 2005; Rusu et al., 2014) whereas our model used a simpler neuron model each with only a single type of excitatory and inhibitory synapse. A recent review by (Ziemann, 2020) stated that there is currently no evidence that I-waves are generated by high-frequency oscillations of individual corticomotoneuronal cells, favouring models of polysynaptic excitatory and inhibitory interneuronal circuits. Although more detailed models could be introduced into our modelling framework, the ability of a simpler population-based model to replicate experimental results suggests that complex neuron-level behaviour may not be essential for the generation of I-waves.

The model used a simplified version of TMS, stimulating a random selection of neurons making up a defined proportion of the population. However, the sites of real TMS stimulation are shown to be dependent on individual neuron properties, such as orientation of the axons relative to the electric field (Rotem and Moses, 2006; Rotem and Moses, 2008; Pashut et al., 2014; Siebner et al., 2022). Previous models of TMS have included computed electrical fields and volume conductor models which take into account complex geometry of the cortex including gyral folding (Seo et al., 2017). Computational models show that the geometry of cortical folding as well as neuronal morphology are key parameters of the effects of TMS (Seo et al., 2017). For the purpose of explaining the origin of I-waves, detailed geometry for stimulation was not taken into account and more complex modelling of the TMS effect on the cortex, for example as an electromagnetic field stimulating axons lying in specific orientations, rather than as direct current inputs to point neurons, could be a future area of exploration.

Future work building on this research could extend the model to include spinal and neuromuscular components to produce motor evoked potentials, further exploring the cortical response and the downstream effects of various experimental TMS protocols. Next steps could also explore the effect of paired pulse stimulation on inhibition, which could suppresses late I-waves in short interval cortical inhibition experiments. It is suggested that sub-threshold TMS mostly activates inhibitory neurons because of their lower threshold for magnetic stimulation (Rusu et al., 2014). Higher stimulation levels might also activate more excitatory neurons or possibly neurons from both AP and PA circuits, contributing a higher number of I-waves being generated as observed in human experimental data (Aberra et al., 2020; Siebner et al., 2022).

Using a spiking neural network model of motor cortical circuits, we have demonstrated how populations of neurons can reproduce observed responses to TMS. The model recreates observations in spontaneous firing, I-wave activity, and responses to increased inhibitory activity in the motor cortex. We also demonstrated how different circuits could contribute to the effects of AP and PA direction TMS, and tested previously hypothesised circuits thought to be involved in the generation of I-waves to show the necessary contribution of more complex connectivity which exists physiologically in the cortex. Spiking neural network models involving large numbers of individual cells are advantageous over morphological models or mean-field models because they enable the observation of contributions of both individual neurons and population-based activity, linking between micro- and meso-scale physiological activity. Developing computational models alongside experimental testing is vital for contributing to the understanding of the effects of TMS and the contribution of cortical circuits to the generation of movement.

## Conclusion

The cortical response to TMS can be explained by recurrent connections in a spiking neural network model of cortical circuitry not tuned to generate oscillatory responses. Parameters that played a significant role in shaping I-wave generation include the refractory period and the transmission delays of neurons. The model explained the generation of I-waves in populations of neurons of the cortex by implementing and testing a range of previously defined circuits. Our results verify that layer 2/3 to layer 5 connections are important in generating I1 and recurrent inhibitory connections play a role in generating I2. More complex circuitry involving other cortical layers was necessary to achieve the generation of I3 as well as realistic resting state activity. With this model, we also proposed a circuit to reproduce the AP response and showed the effects of specific model parameters and increased inhibitory activity. This work extends previous studies of cortical modelling to understand experimentally measured behaviour in the human nervous system.

## Acknowledgments

The author thanks John Cirillo for their valuable discussions and Mark Sagar for initial project funding and administration. This preprint was formatted using the LaPreprint template (https://github.com/roaldarbol/lapreprint) by Mikkel Roald-Arbøl.

## Author contributions

Conceptualisation: A.M., L.H.; Methodology: A.M., L.H.; Software: L.H.; Writing original draft: L.H.; Review & editing: A.M., T.B., L.H.; Supervision: A.M. T.B.; Administration & Funding acquisition: A.M., T.B.

